# Statelets: High dimensional predominant shapes in dynamic functional network connectivity

**DOI:** 10.1101/2020.08.16.252999

**Authors:** Md Abdur Rahaman, Eswar Damaraju, Debbrata Kumar Saha, Sergey M. Plis, Vince D. Calhoun

## Abstract

Dynamic functional network connectivity (dFNC) analysis is a widely used approach for capturing brain activation patterns, connectivity states, and network organization. However, a typical sliding window plus clustering (SWC) approaches for analyzing dFNC continuously models the system through a fixed set of connectivity patterns or states. It assumes these patterns are span throughout the brain, but in practice, they are more spatially constrained and temporally short-lived. Thus, SWC is not designed to capture transient dynamic changes nor heterogeneity across subjects/time. Here, we adapt time series motifs to model the temporal dynamics of functional connectivity. We propose a state-space data mining approach that combines a probabilistic pattern summarization framework called ‘Statelets’ — a subset of high dimensional state-shape prototypes capturing the dynamics. We handle scale differences using the earth mover distance and utilize kernel density estimation to build a probability density profile for local motifs. We apply the framework to study dFNC collected from patients with schizophrenia (SZ) and healthy control (HC). Results demonstrate SZ subjects exhibit reduced modularity in their brain network organization relative to HC. These statelets in the HC group show more recurrence across the dFNC time-course compared to the SZ. An analysis of the consistency of the connections across time reveals significant differences within visual, sensorimotor, and default mode regions where HC subjects show higher consistency than SZ. The introduced statelet-approach also enables the handling of dynamic information in cross-modal applications to study healthy and disordered brains and multi-modal fusion within a single dataset.

## 1. Introduction

Dynamic functional network connectivity (dFNC) enables the investigation of temporal changes in connections among different brain regions. It has been established as a powerful technique for analyzing time-varying activity coordination across brain regions (Sakoğlu, Pearlson et al. 2010; Allen, Damaraju et al. 2014). The Pearson correlation typically characterizes the coordination, computed using a sliding window (e.g., 44s in length) technique for measuring the interdependence among brain regions or independent components of interest extracted from group ICA (Allen, Damaraju et al. 2014; Damaraju, Allen et al. 2014). Recent empirical evidence has strongly suggested that this dynamic spatiotemporal configuration better models brain activities and network organization (Sakoğlu, Pearlson et al. 2010; Ma, Calhoun et al. 2014; Liu, Wang et al. 2017; Vergara, Mayer et al. 2018; Zhi, Calhoun et al. 2018). dFNC analysis provides an ability to track time-varying transitions in connectivity strengths, thus moving beyond the firm and apparently incorrect assumption of a single static connectivity pattern (Damaraju, Allen et al. 2014; Rashid, Damaraju et al. 2014; Vergara, Mayer et al. 2018). The kinetics are shown to be informative about the temporal variations in brain network organizations - which have already been proven as a dynamical system (McKenna, McMullen et al. 1994; Bullmore and Sporns 2009). A typical dFNC analysis assumes fixed discrete states with varying occupancy over time and does not fully capture recurring transient information. A few studies analyze dFNC considering brain transitions through states of connectivity and estimate those states via clustering the windowed FNC sliding across time (Damaraju, Allen et al. 2014; Miller, Yaesoubi et al. 2016; Saha, Abrol et al. 2019). However, the obtained states essentially fuse all homogenous connectivity patterns ignoring their order and timing. For a given dFNC time course, the methods do not distinguish between predictable versus unexpected patterns and consider all time-steps equally significant. These states explore connectivity patterns that span throughout the brain and prevail for a more extended period. In practice, connectivity signatures are more spatially constrained and short-lived (Miller, Yaesoubi et al. 2016). That is, transformations modulate over a slower timescale, and dynamics can adjust functional network topology accordingly (Bullmore and Sporns 2009; Li, Zhang et al. 2017). So, these methods are less suitable for capturing brief dynamic changes in the dFNC time course; consequently, they miss the dynamic properties of dFNC, such as network topology, synchronizability, and intermittent connectivity, which are affected by neuropsychiatric disorders like schizophrenia (Yu, Allen et al. 2012; Siebenhühner, Weiss et al. 2013). Thus, in this paper, we address the tradeoff between the desire to model transient dynamics and the need to constrain the set of extracted features to a reasonable size.

Modern neuroimaging studies focus on data-driven symptom investigation and bio-marker discovery, which depend on identifying interpretable subsets of data that characterize diseases and controls (Nielsen, Hansen et al. 2004; Atluri, Padmanabhan et al. 2013; Kong and Yu 2014). In data mining, time series analysis has been receiving significant attention in the past decades. Recent studies have established ignoring some data contribute to better data mining (e.g., clustering/subgrouping) (Rakthanmanon, Keogh et al. 2011). Consequently, time series motifs as such signatures are powerful tools for modeling and analyzing dynamical systems (Mueen 2014; Torkamani and Lohweg 2017). The motif is defined as a previously unknown and the most recurring pattern of a time series (Lin, Keogh et al. 2002). In recent years, motifs discovery and analysis have become a popular medium for observing these intermittent events (Lin, Keogh et al. 2002; Chiu, Keogh et al. 2003; Wilson, Birkin et al. 2008; Mueen, Keogh et al. 2009; Gao and Lin 2018). In a similar spirit, we propose a state-space data mining approach called “Statelets”. We implement earth mover distance (EMD): a simple yet effective similarity metric for motif comparison. EMD provides a scale-independent comparison between signatures, can handle variable-length substructures, and accounts for partial matching. EMD is successfully applied as a distance metric for comparing distributions in image retrieval (Rubner, Tomasi et al. 2000). Motifs extraction from individual time series often yields an enormous number of variable-length shapes for a large dataset. The superfluousness of data points limits the model’s ability to generalize the trends, thus predicting the overall dynamical system’s behaviors is still looking for a needle in a haystack. Therefore, summarizing local motifs have become cardinal for a comprehensive dynamic investigation. Studies that proposed summarizing time series are mostly domain-specific and suffer from a lack of generalization (Sripada, Reiter et al. 2003; Ahmad, Taskaya-Temizel et al. 2004; Kacprzyk, Wilbik et al. 2008). In this paper, driven by the desire to build a more general summarization method and equipped with an efficient EMD implementation, supporting kernel density estimation (KDE) (Silverman 1986) in motif space, we propose a novel probabilistic pattern summarization framework called ‘statelets. Obtaining descriptive statelets from patients and controls helps interpret the group differences and explainable features from both trajectories.

A previous study suggests that tracking down the representative patterns from the dFNC time series enhances the sensitivity of the outcomes and provides a more accurate assessment of the dynamic (Morioka, Calhoun et al. 2020). We explore the dynamic behavior of a healthy and schizophrenic brain and their group differences. The probability density of group statelets reveals unique sequences of connections among functional brain networks (Fig. 9) that provide new insights into the functional connectivity changes associated with the disorder. Results also evident reduced modularity in patient’s (SZ) dynamics relative to healthy controls (HC) (Fig. 11). Passage coding – convolving the corresponding statelet with the time course of both dynamics establishes patterns in HC that are more frequently recurring than in SZ, reflected in the time decay graph (Fig. 12). Furthermore, the experiment of the transitivity of time decay graphs indicates that HC networks intercommunicate more and consistently keep the channels active compared to SZ (Fig. 13). These experimental results on the human brain’s network organization’s dynamic features provide a better explanation of multiple symptoms of schizophrenia, e.g., disorganized thinking, inefficient information processing, and decision-making. Consistency of the connections across time reveals significant differences within visual (VIS), sensorimotor (SM), and default mode (DM) regions where HC subjects show higher consistency than SZ (Fig. 15). Finally, statelets wise subgrouping of the dynamics report salient and stable group differences in sensorimotor (SM), cognitive control (CC), and cerebellar (CB) domains, which are not observed in earlier studies (Fig. 17).

## 2. Definitions and BackGround

### A. Definition 1: Dynamic functional network connectivity

Dynamic functional network connectivity (dFNC) is a time-varying correlation typically computed using a sliding window (block of time points, e.g., 30-60s) technique among independent components (ICs) of the brain. The correlation value at each window is also a quantification of the functional connectivity strength between a pair of brain networks within the time frame.

### B. Definition 2: dFNC Time Course

A dFNC time series, T = W1, W2, W3, ......, Wn is a sequential set of n real values, where Wi represents the correlation between two independent components (ICs) of the brain for a certain period. W stands for the window.

### C. Definition 3: Connections/pair

A connection is a functional association between a pair of independent neural components. We used ‘connection’ and ‘pair’ interchangeably in our writing.

### D. Definition 4: Subsequence

Given a time series *T* of length *l*, a subsequence *S* is a subset of length *m ≤ l* contiguous indices of *T*.

### E. Definitions 5: Motif

Given a time series *T*, a motif *H*_*i, m*_ is a subsequence of *T* having the minimum average EMD distance from *k* other subsequences of *T* with length *m*. A module for finding local minima selects *k* to determine no-overlapped matches. That is, *H*_*i, m*_ is the most recurring shape in *T* of length *m*.

### F. Definition 6: Statelets

Given a collection (constant/variable-lengths) of time series motifs *G* and a positive real number *d* (size parameter), Statelets *S* is a subset of size *d* of state-shape prototypes evaluated using the proposed summarization framework.

Functional magnetic resonance imaging (fMRI) of the human brain has been used broadly for studying brain disorders (Gur, McGrath et al. 2002; Calhoun, Adalı et al. 2006; Swanson, Eichele et al. 2011). Outcomes from fMRI analysis reinforce the intuition of dysconnectivity in disorders such as schizophrenia, i.e., unusual connections among distinct brain networks (Stephan, Baldeweg et al. 2006; Williamson and Allman 2012; Damaraju, Allen et al. 2014). The decomposition of neural features and generating biomarkers helps distinguish schizophrenia to a greater extent (Calhoun, Liu et al. 2009; Erhardt, Rachakonda et al. 2011; Du, Fu et al. 2019; Rahaman, Turner et al. 2019). Nevertheless, the generalization of the outcomes and being consistent in exploring the neural system remains challenging because of the heterogeneous nature of neuropsychiatric disorders such as schizophrenia (Tsuang, Lyons et al. 1990; Alnæs, Kaufmann et al. 2019). Thus, studies employ subgrouping/clustering of the subjects to minimize the dissimilarity and making a more rational comparison (Luchins, Weinberger et al. 1979; Scarr, Cowie et al. 2009; Rahaman, Damaraju et al. 2020). A typical sliding window plus clustering (SWC) analysis approach resolves the problems by continuously modeling the system through a fixed set of connectivity patterns or states (Allen, Damaraju et al. 2014; Damaraju, Allen et al. 2014; Miller, Yaesoubi et al. 2016; Vergara, Mayer et al. 2018; Saha, Abrol et al. 2019). Moreover, a recent study of tri-clustering dFNC data suggests three-dimensional connectivity states and reveals more significant group differences than previous studies (Rahaman, Damaraju et al. 2020). However, the most widely used SWC mechanisms assume each window carries information from all the pairs (connections); consequently, the approach explores patterns that span throughout the brain. In practice, connectivity signatures often spread over a smaller subset of components (spatially), and different subsets showcase different types of connectivity patterns. Also, the patterns manifest across a smaller number of windows (temporally). Hence, we need an approach that allows us to investigate the system more independently. That is, a way to evaluate the time series of each pair individually and identify potential short length interactions between two components and their geometric structures (shape). Another deficiency of this method is that there is no accounting for the recurrence of a signal pattern. Since dFNC reflects the interaction between a pair of brain components, we are more interested in repetitive patterns across the timeline. We can adapt this by extracting motifs of the time course as patterns occurring very sparsely might be noise or artifacts in our case.

Exploring the entire time series at a time yields an inefficient way to exploit the relations. As an example, we can show the following scenario where we take two random dFNC time courses. Figure 1 shows the conventional way of comparing the time series in the previously reported dFNC state analysis studies. Time courses A & B look different if we consider the whole sequence. Considering all the data at once makes it harder to observe the similarities in these signals. Hypothetically, more sparse patterns are unlikely to represent the interaction between neural components, thus less helpful to learn the dynamics. Instead of generating some fixed-length signatures with limited temporal information, statelets account for the dynamic changes in functional brain connectivity and precisely quantify the patterns leading the dynamics, including their temporal and spatial adjustments. We can see that the comparison has improved drastically in figure 2. Both time series look very similar to each other using the dominant patterns only. In the earlier comparison, the differences create redundant states that potentially complicate the outcomes. Creating more states may help with that issue, but adding more states can also result in assigning a lot of loosely related windows to each of the *k* states since researchers often use *k*-means clustering (Li, Zhang et al. 2017; Zhang, Zhou et al. 2019) for estimating dFNC states.

**Fig. 1.**
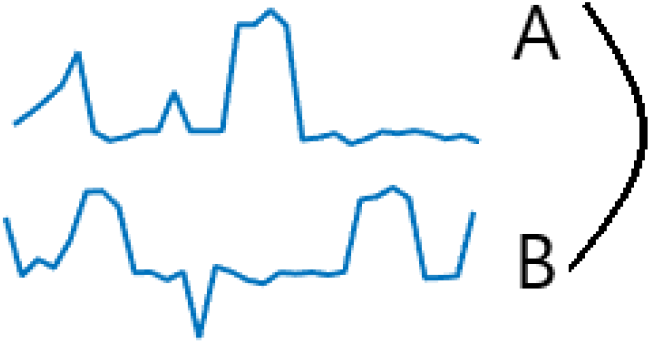
Comparison between two arbitrary signatures A and B in a conventional clustering method. It considers the whole sequence for measuring the distance (e.g., Euclidean). In the dFNC time course, it considers a connectivity pattern for a given window W_i_ across all the connections. For each window, the length of the signature is n, where n is the number of pairs of components.

**Fig. 2.**
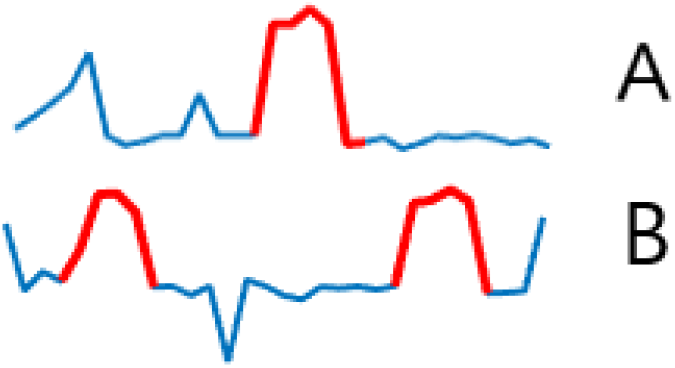
Comparing only the motifs (bold red) of dFNC time courses.

Furthermore, model order selection is a free choice in state-of-the-art methods. We tried to address it in our summarization framework by capturing state shapes representation of all intrinsic subgroups. To our best knowledge, this is the first endeavor to outline the state shapes from local motifs based on their probability density throughout the datasets.

## 3. Data collection and Preprocessing

We used the fBIRN dataset for generating the dFNC time courses in this project (Keator, van Erp et al. 2016). The data repository has resting-state functional magnetic resonance imaging data collected from 163 healthy controls (117 males, 46 females; mean age 36.9), and 151 age-and-gender matched patients with SZ (114 males, 37 females; mean age 37.8) during the eyes-closed condition. Collected data pass-through data quality control (explained in Damaraju et al., 2014 (Allen, Damaraju et al. 2014)). The participant’s consent was obtained before scanning following the Internal Review Boards of affiliated institutions. Data were collected with a TR of 2 seconds on 3T scanners. Imaging data for six of the seven sites were collected on a 3T Siemens Tim Trio System and a 3T General Electric Discovery MR750 scanner at one site. Resting-state fMRI scans were acquired using a standard gradient-echo echo-planar imaging paradigm: FOV of 220 × 220 mm (64 × 64 matrices), TR = 2 s, TE = 30 ms, FA = 770, 162 volumes, 32 sequential ascending axial slices of 4 mm thickness and 1 mm skip. Subjects had their eyes closed during the resting state scan. Data pre-processing, quality control, and dFNC collection follow the standard pipeline described in (Damaraju, Allen et al. 2014). Firstly, rigid body motion correction has been done using the INRIAlign toolbox in SPM to correct for subject head motion, followed by a slice-timing correction to account for timing differences in slice acquisition (Freire and Mangin 2001). Then the scans went through a 3dDespike algorithm to regress out the outlier effect and warped to a Montreal Neurological Institute (MNI) template and resampled to 3 mm^3^ isotropic voxels. Instead of Gaussian smoothing, we smoothed the data to 6 mm full width at half maximum (FWHM) using the BlurToFWHM algorithm, which performs smoothing by a conservative finite difference approximation to the diffusion equation. Additionally, The voxel time course was variance normalized before perform independent component analysis (ICA) (Hyvärinen and Oja 2000) as this has shown better to decompose subcortical sources in addition to cortical networks. Then, group ICA (Calhoun, Adali et al. 2001) was performed on the pre-processed data and identified 47 intrinsic connectivity networks (ICNs) from the decomposition of 100 components. Subject-specific spatial maps (SMs) and time courses (TCs) were obtained using the spatiotemporal regression back reconstruction approach implemented in GIFT software (Calhoun, Adali et al. 2001). After the ICA, we obtained one sample *t*-test maps for each SM across all subjects and threshold these maps to obtain regions of peak activation clusters for that component; we also computed the mean power spectra of the corresponding TCs. These heuristics curated a group of components as intrinsic connectivity networks (ICNs) if their peak activation clusters fell on gray matter and showed less overlap with known vascular, susceptibility, ventricular, and edge regions corresponding to head motion. The mean sFNC matrix was computed over subjects. The 47 components are organized into modular partitions using the Louvain algorithm of the brain connectivity toolbox (dFNC between two ICA time courses was evaluated using a sliding window approach with a window size of 22 TR (44 s) in steps of 1 TR (Calhoun, Adali et al. 2001; Allen, Damaraju et al. 2014). The window constituted a rectangular window of 22 time points convolved with Gaussian of sigma 3 TRs to obtain tapering along the edges (Allen, Damaraju et al. 2014). We estimated covariance from the regularized inverse covariance matrix (ICOV) (Varoquaux, Gramfort et al. 2010; Smith, Miller et al. 2011) using the graphical LASSO framework (Friedman, Hastie et al. 2008). The regularization parameter was optimized for each subject by evaluating the log-likelihood of the subject’s unseen data in a cross-validation framework—all post hoc steps for extracting and validating. The subject wise dFNC values were Fisher-Z transformed and residualized with respect to age, gender, and site.

## 4. Earth Mover Distance (EMD)

We implement the Earthmover distance (EMD) for computing distances between motifs. The EMD is a measure of dissimilarity between two probability distributions (Vallender 1974; Levina and Bickel 2001). It corresponds to the minimum amount of work required to make two distributions look-alike (Champion, De Pascale et al. 2008). For *p* and *q* two given distributions, *F (p, q)* represents the set of all possible flow between *p* and *q*. Then, the work is defined as follows,

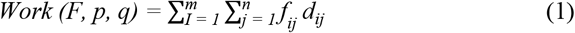

here *d*_*ij*_ is the distance between *p*_*i*_ and *q*_*j*_, m and n denote the number of clusters in distribution *p* and *q*, respectively. The EMD between *p* and *q* is the minimum amount of work required to match these two distributions given by the following equation (Andoni, Indyk et al. 2008; Champion, De Pascale et al. 2008).

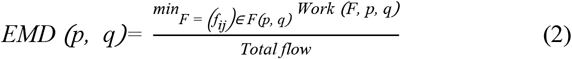

### Implementation in our study

We use a particular case of EMD for a one-dimensional time series, which parses through the vectors and keeps track of how much flow must be done between consecutive time points. Here, flow is the difference between the amplitudes of two shapes at a given time point. The method recursively accumulates the absolute work done at each time point, and finally, the summation over the time points corresponds to the distance (see equation 5 and 6). Equation 3 and 4 normalize shapes P and Q, respectively.

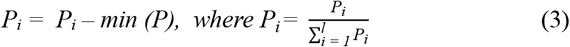

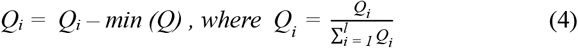

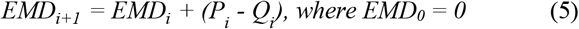

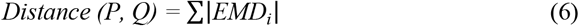

## 5. Our proposed framework

Given this time series dataset, our architecture performs two fundamental steps to estimate state-shape prototypes (statelets), A) extraction of local time series motifs, and B) summarization using associated probability density. Figures 3 depicts the subprocesses required to perform these steps. The time-varying correlation between two resting-state network maps is computed using a sliding window technique described in section 3. This step produces dFNC time courses we use to discover the motifs.

**Fig. 3.**
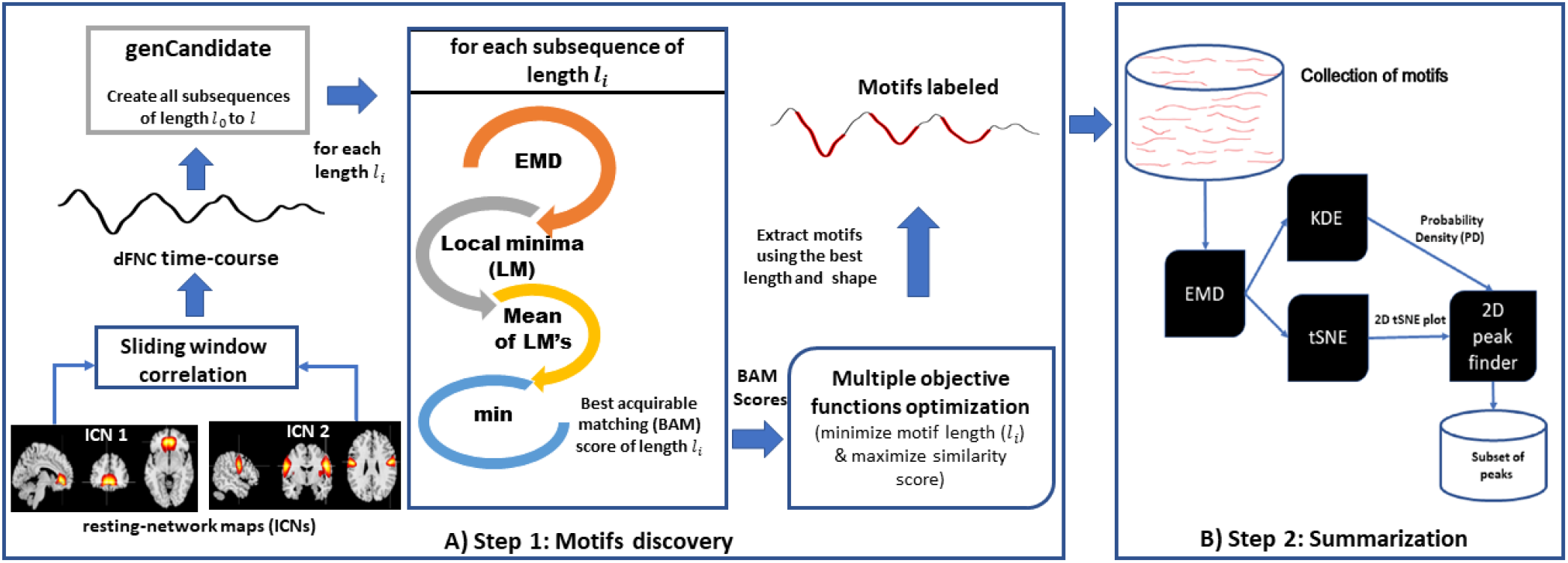
Our proposed methodology for time series motifs discovery and summarization. A) Step 1: Extraction using EMD as a similarity metric. For a given time course, the subroutine extracts the most repetitive pattern along with their multiple occurrences. B) Step 2: Probabilistic summarization of local motifs using KDE. It approximates a subset of the most frequent shapes/patterns covering all the subgroups from a given collection of varying length motifs. The distance matrix computed using EMD is used to plot tSNE and for computing probability density (PD).

### A. Step 1: Local motifs discovery

Each dFNC time series represents the functional connectivity between two distinct networks of the brain. Therefore, it is not feasible to impose a common motif length for all the time series in the dataset, like most other motif discovery methods. The proposed heuristic takes a range [R1 R2] and suggests the perfect length to explore the time series within that boundary. In our case, dFNC is collected using a window size of 22 time points. So, the lower bound for R1 > 22. The upper bound for R2 should be confined within half of the signal length for a sensible parcellation of dFNC. Since the study aims to observe transient signatures in the data, we select the range from 30 to 50 in our analysis. At first, the process creates all the subsequences of different lengths and then passes these candidates through the following subroutines. Step1 in figure 3 (A) demonstrates the method for performing local motifs discovery. At first, the process creates all the candidate motifs of different lengths and passes them through the following mini layers.

#### 1) EMD layer

For a given candidate of length *l*_*i*_, the algorithm searches through the timepoints by taking all other candidates of that length and computing the EMD between them. We can consider the selected candidate as a kernel here, and the idea is to apply that kernel on the input signal to compute distances.

#### 2) Pooling Local Minima (LM)

The module pools a subset of top matches (minimum distance) instead of just one across the signal. Since motifs are repeating signatures, intuitively, we scrutinize the top matches only. It divides the input time course into several subsections and pool one minimum from each subsection.

#### 3) Mean of LM’s and global minimum

This layer computes the mean of those local minima, representing the overall performance by that specific candidate. After finishing computation for all the candidates of *l*_*i*_, the method selects the global minimum, which corresponds to the best acquirable matching score (BAM).

#### 4) The best length selection

To extract most recurring transient patterns, a recommender unit optimizes two objectives on the BAM scores from all the lengths, minimizing the motif’s length and maximizing the similarity score to select the best length.

#### 5) Motifs extraction

In the length selection process, we have already collected the information required for motifs extraction. So, using the leading candidate as the benchmark, the subroutine collects similar non-overlapping occurrences of local motifs across the time course and store these motifs for creating global dominants across the subjects for a given pair.

### B. Step 2: Sumamrization

For each pair of components (connection), we discover a group of shapes from all the subjects. Now, we run a module called “FindDominants” consist of the four steps (a, b, c, and d) mentioned in figure 3 (B) to identify pairwise dominant motifs. These shapes cover all the subjects and pairs of the group (SZ/HC). We estimate the dominants across the dataset using the probability density of the motifs in our collection. To do this, we extract the representative of each subgroup, which we call statelets. Each statelet has a persistence level, which is given by their probability density across group dynamics. We follow the same strategy for both subject groups SZ and HC.

#### a) Distance measure

We use EMD as the distance measure after normalizing the time courses to make our distance invariant to scale and offset. Then, compute cross EMD across the shapes, which creates a square matrix of *n* × *n* dimension where *n* is the number of shapes in the given collection.

#### b) Kernel density estimator (KDE)

Our method applies KDE to calculate probability density by each of the motifs in the subset. We incorporate KDE with a Gaussian kernel and optimal bandwidth (Silverman 1986; Sheather and Jones 1991). The kernel density at a point × is given by equation (7),

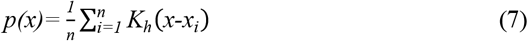

For ‘*D*’ dimensions, the formulation becomes,

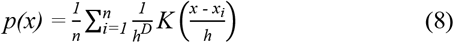

where *n* is the number of samples, *k*_*h*_ is the kernel with a smoothing parameter h called the bandwidth. The kernel is a non-negative function and *h* > 0. For a smoother density model, the common practice uses the Gaussian kernel, simplifying the following kernel density model.

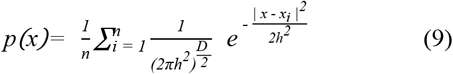

here *h* is the standard deviation of the Gaussian components. Another crucial part of the process is to select an appropriate bandwidth (*h*) for the density, and there are several strategies for selecting *h* (Turlach 1993; Ahmad and Amezziane 2007). Our approach incorporates the rule-of-thumb bandwidth estimator for Gaussian (Scott 2012). The optimal choice for *h* is following,

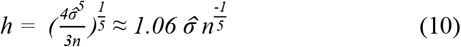

here 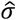 is the standard deviation of the samples, and *n* is the number of total samples. The term (*x* - *x*_*i*_) in the equation corresponds to the distance between the sample *x* and *x*_*i* ∈ *n*_. So, we assign EMD distances (previously computed) between the shapes as a value of (*x* - *x*_*i*_) in the equation. Using equation (9), KDE generates probability density for all motifs in the collection.

#### c) tSNE using EMD score

t-distributed stochastic neighbor embedding (tSNE) leads to a powerful and flexible visualization of high-dimensional data by giving each datapoint a location in a two or three-dimensional map (Maaten and Hinton 2008; Gisbrecht, Schulz et al. 2015). Dcentralized stochastic neighbor embedding (dSNE) is used to separate the data subgroups using their distance metrics (Saha, Calhoun et al. 2017; Saha, Calhoun et al. 2019). tSNE considers each shape a data point in the subspace and uses EMD distances to select the neighbors for the embeddings. Then, the method weighs all the points using their corresponding probability density. Figure 4 shows all possible subgroups revealed by tSNE. The color indicates the probability density. After that, the method maps all tSNE points to a two-dimensional grid to accumulate the probability density of all closely neighboring shapes onto a single cell. It creates an erratic increment of frequency in the corresponding vicinity, making the peak points more divergent. The process discretizes the range of coordinates into a set of integer intervals. As a result, multiple points from the tSNE plot are stacked together into one cell (figure 5).

**Fig. 4.**
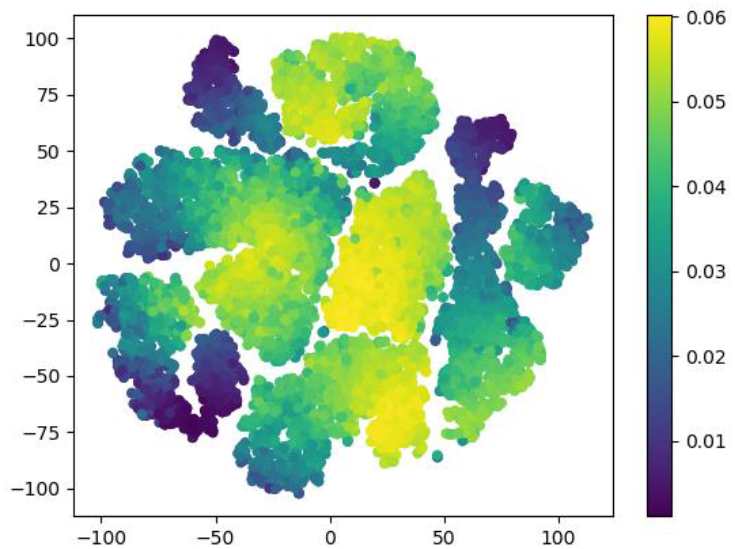
tSNE using EMD distance. Data points represent the motifs weighted by their probability density computed using KDE. X and Y-axis stand for the horizontal and vertical coordinates of each point, respectively.

**Fig. 5:**
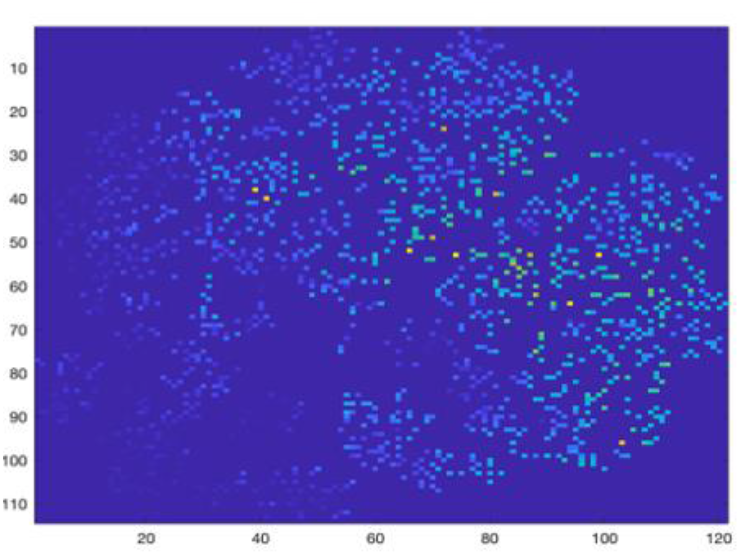
Mapping tSNE points to a 2D matrix for accumulating frequency of close neighbors. Therefore, we observe the higher density data with a brighter color in the figure.

#### d) Gaussian Blur and peak finding

Defocusing noise helps extract significant features/data points representing the subgroup peaks.

Subsequently, we blur the tSNE image (figure 6) and use a two-dimensional peak finder to compute the high-density data points. This extracts a characteristic shape from each subgroup with a higher probability density (PD) (figure 7). For more clarity, the method restricts the peak finding to a certain level to not end with a subset of similar shapes. The objective is to find a diversified set of shapes which well approximates the overall dynamics. These peaks are considered as statelets for that collection.

**Fig. 6.**
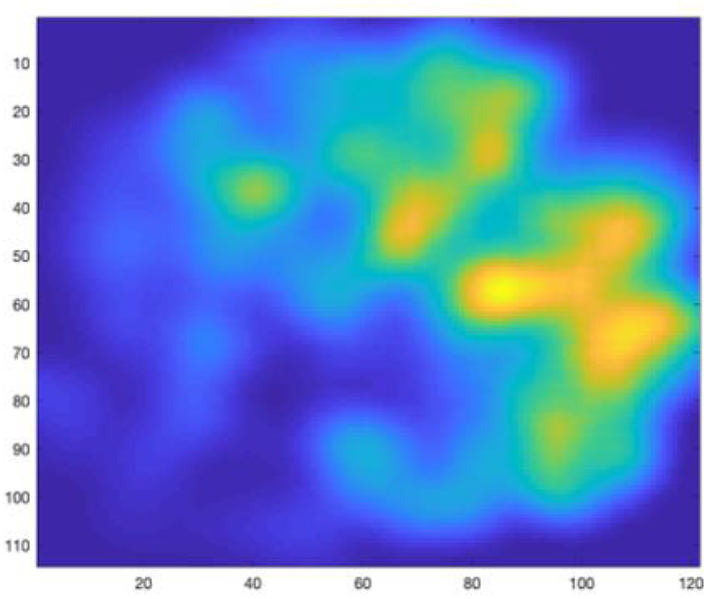
After applying the Gaussian Blur on the 2D image to defocus less dense data points.

**Fig. 7.**
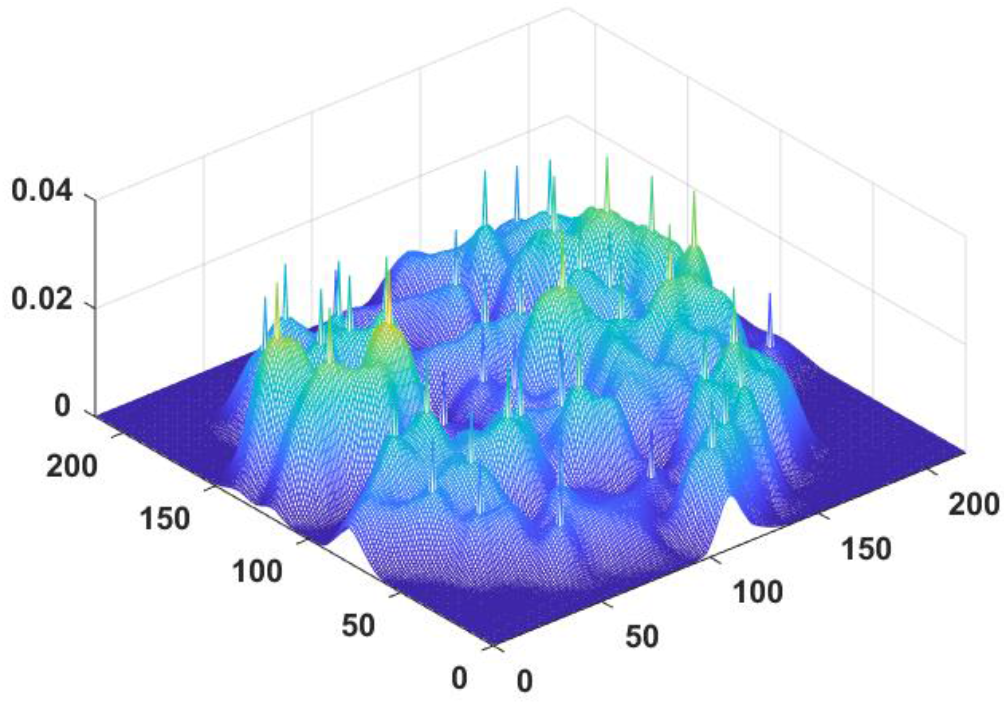
A three-dimensional view of the tSNE plot after marking the peaks (extracted by 2D peak finder) from each subgroup. The peak finder selects at least one peak from each high-density region. Later, we map these peaks tSNE coordinates back to determine real motifs the points representing.

## 6. Experimental Results

We divide the dataset into two groups of 163 healthy control subjects (HC) and 151 schizophrenia patients (SZ). Each subject has ^47^*C*_2_ = 1081 pairs of components: thus, 1081 dFNC time courses. For each pair, we take time courses from all the subjects within that group (HC/SZ). We have 151 and 163 dFNC time courses per pair of components for SZ and HC, respectively. Our method evaluates a subset of statelets from both groups of subjects (SZ/HC). The probability density of these high dimensional state shapes denotes the replicability of those statelets across connections among multiple brain regions. In figure 8, we show a subset of the most frequent statelets from both groups. In figure 9, each bar indicates how often a pair of components has appeared in the group dynamics. They are also sorted based on their relative rank computed on their frequency in both patient and control groups.

**Fig. 8.**
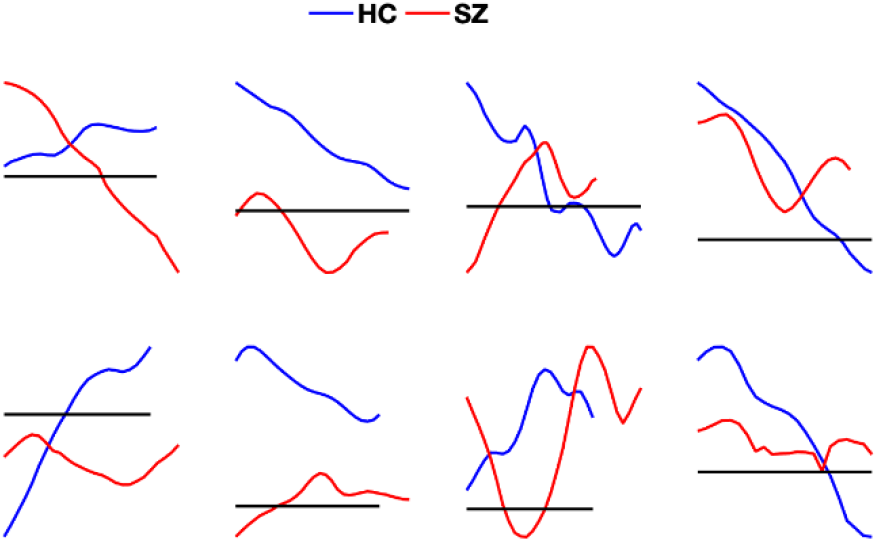
A subset of dominant statelets from SZ and HC dynamics. Each subplot shows two shapes from both phenotypes sorted according to their overall probability density. The red and blue correspond to patient and control groups, respectively.

**Fig. 9.**
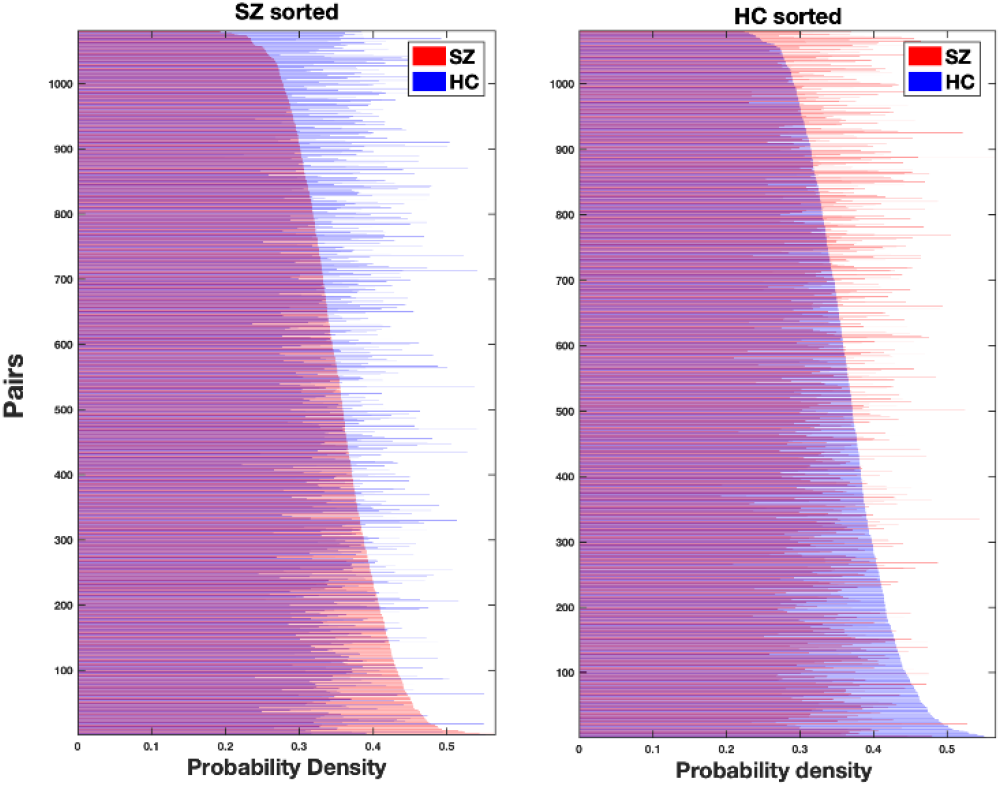
The probability density (PD) of the connections demonstrates how frequent the statelets extracted from a connection appear in the entire group dynamic. Each connection has two density values – one per subject group. Blue demonstrates a probability density in controls (HC) and red in patients (SZ). In the left subplot, we sort the connections low to high according to their probability density in the SZ group, and HC sorts the right subplot. Y-axis represents the order of the links after sorting, and X-axis depicts their corresponding probability density. We observe that the rank of connections differs in both dynamics in terms of their PD, and we show this difference is statistically significant by hypothesis testing.

We can see that the pair (connections) rank is very different in each group, suggesting a diverse influence of connections driving the dynamics of patients vs. controls. The frequency of statelets in each connection in HC is higher than the SZ. To test the statistical significance of this difference between the two distributions, we computed a Kendall tau rank correlation coefficient, which is traditionally used to measure the ordinal association between two entities (Kendall 1938; Puka 2011). The hypothesis test evaluates τ = 0.045 with a p-value = 0.025 indicating a statistically significant difference.

### Dynamic features analysis

We focus more on the dynamic features of the neural system, which have been mostly not measurable in previous investigations of dFNC.

#### a) Recurrence

In figure 10, we show the occurrences of the group statelets in individual dynamics (randomly selected three subjects from each group).

**Fig. 10:**
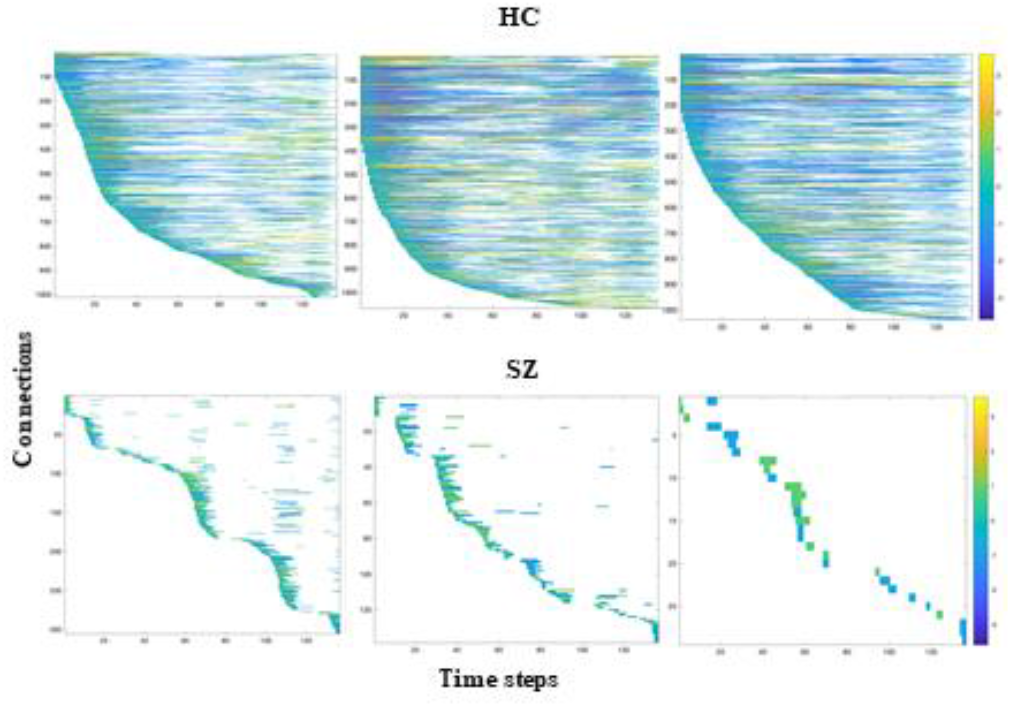
Each graph represents the occurrences of the most recurring statelets over each subject’s time course. These are three randomly selected subjects from the HC (top) and SZ (bottom) group. We convolve the group statelets with the subject dynamics (all the dFNC time courses, 1081) to investigate the recurrence of the statelets over time. Then, we sorted the pairs based on the dominant shape’s first occurrence in their time series. Consequently, the early the statelets appear in a pair’s time course, the higher the pair/connection in that subject’s dynamics—we threshold these convolutions matching scores at 0.8 for both groups to track down the strong appearances only. The color intensity corresponds to how strongly/weakly the shape appears in that part of the course. The color bar is identical for all the reference subjects in the figure.

We investigated the recurrence of statelets for all the subjects and got consistent results, as shown in figure 10. Statelets are the most generalized envelopes of connectivity. dFNC sequences from HC groups converge to these envelopes more often across the timeline. The higher recurrence endorses better synchronization in the brain circuitry. In the SZ dynamic, the number of repetitions of statelets is significantly lower, which initiates the aberration and inefficiently controls the flow of information. This might be responsible for the cognitive symptoms like difficulty in processing information to make decisions. Also, a smaller number of connections are present in the SZ dynamics than HC. SZ group statelets characterize a lower number of pairs, which indicates more chaotic connections in SZ dynamic. This graph also demonstrates the temporal modifications in statelets of a connection. The higher convolution value (brighter color) indicates the steady occurrence of the statelets, i.e., with similar shapes, but the lesser values stand for the distorted appearance. So, it explains the gradual modification in the state shapes with time and clarifies the rebounding to the earlier forms.

#### b) Modularity

The line graphs in figure11 illustrate the ensemble behavior of the connections between a pair of networks in both dynamics (SZ/HC). At each time point, the value represents the number of connections with similar connectivity patterns. Throughout the scan session, comparatively more HC connections are activated in identical time steps than patients. This highlights the higher modularity in the HC network/graph (the connections between the functional components of the brain). Modularity is imperative because it quantifies the strength of the network’s division into modules, and each module consists of a subset of nodes, which are independent brain components in our case. These modules help data transmission by performing specific tasks independently. The study already established individuals with higher baseline modularity exhibited greater improvements with cognitive training, suggesting that a more modular baseline network state may contribute to greater adaptation in response to cognitive training (Arnemann, Chen et al. 2015). Also, the cognitive ability is influenced by the relative extent of integration and segregation of functional networks (i.e., modules) distributed across distant brain regions (Chen, Abrams et al. 2006; Stevens, Tappon et al. 2012). Subsequently, the difference in modularity is informative about the cognitive symptoms of schizophrenia. In addition, the graph shows SZ connections become modularized at the end of the time series, whereas the HC ensembles are reduced by their size (figure 11). This ensemble activation also demonstrates a distinct synchronization pattern between SZ and HC brain and suggests an interchanging control by various neural components.

**Fig. 11.**
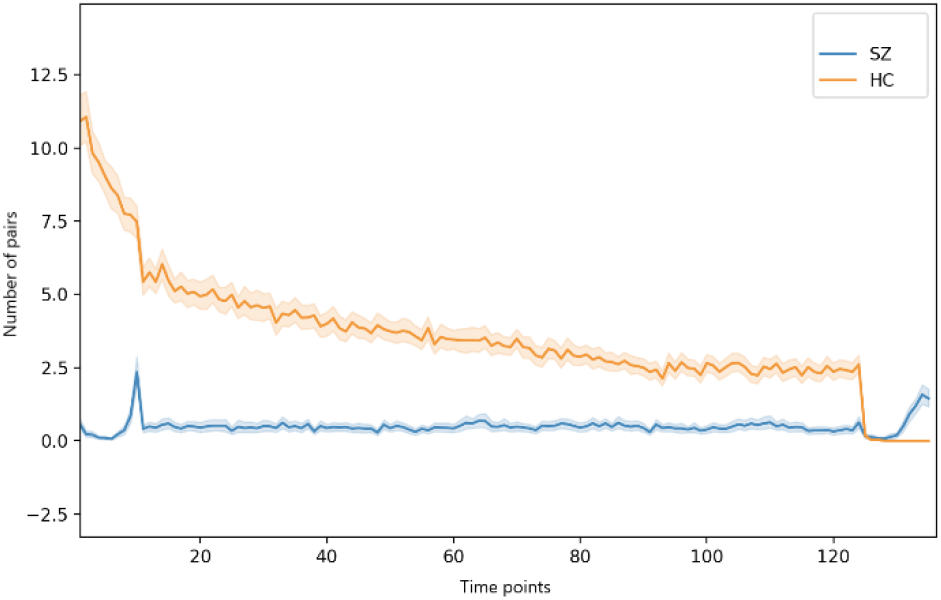
Based on probability density ranking (in figure 9), we computed a collective appearance of the connections across all the subjects. The X-axis shows the time steps, and the Y-axis corresponds to the number of pairs connections that show the first statelet at that step, which indicates the activation of the pair.

#### c) Consistency graph

For measuring the consistency of the connections, we introduce a metric called time decay defined as follows.

##### Time decay (TD): a consistency measure

It approximates how consistently a pair’s connectivity exhibits the dominant motif early in the time course. The following equation depicts the idea of *TD*.

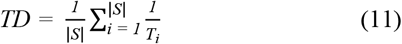

Here, |*S*| = The total number of subjects in the group SZ/HC and *T*_*i*_ the index at which the pair has appeared for the first time in a subject (*T*_*i*_ = 1 to 136). If the pair is not present in a subject’s dynamic, then *T*_*i*_ = ∞ thus, time decay for that pair in a subject is zero. TD can characterize a connection as consistently late (low TD value), consistently early (high TD value), pseudo-random positioning (medium TD value e.g., 0.10 to 0.15), consistent but sparse in the group dynamics (0.5 – 0.09), etc.

Figure 12 demonstrates the average time decay graphs computed on both dynamics. Here, the vertices are independent brain networks, and the edge represents the connection between two functional networks. The weight of the edge stands for the time decay; the darker, the higher. As we can see, HC functional networks are firmly connected, and most of the connections (out of 1081) pass the time decay benchmark. On the other hand, only a small subset of SZ connections converge to their group statelets. That evident that the consistency of SZ network connections is considerably lower than HC. The functional connection between different brain components shows transient and sporadic patterns in SZ; consequently, the information processing experiences aberrant neural pathways. Overall decision making and immediate response towards an action become slower than a control’s brain. The lack of steadiness in channels portrays an inefficient neural signal processing unit that affects multiple cognitive behaviors like organized thinking, trivial reactions, focusing or paying attention, using information immediately after learning it, etc. In figure13, we show the histograms of transitivity of subject wise time decay (TD) graphs in both groups. Transitivity symbolizes the global probability of the network (connectivity graph) to have adjacent nodes interconnected, revealing the existence of tightly connected communities, e.g., clusters and subgroups. The majority of the SZ graphs have zero or very low transitivity compared to the HC. We run a two-sample t-test on the transitivity scores from both groups, and the p-value, *p* = 9.23e-88, indicates the difference is statistically significant at a 1 % significance level. The higher transitivity manifests the presence of modules in the network that has been shown to be useful to better cognitive performance (Arnemann, Chen et al. 2015). The analysis indicates that HC networks intercommunicate more consistently and keep the channels up compared to SZ. This result is consistent with previous work (Karlsgodt, Sun et al. 2010; Allen, Damaraju et al. 2014; Damaraju, Allen et al. 2014), which suggested that the patients exhibit more erratic and less efficient communication among brain regions than controls. Figure 14 shows the average time decay of each connection in both SZ and HC groups. The rightmost map shows HC-SZ group differences. All the pairs in HC show a more substantial time decay that SZ, especially in visual, sensorimotor, and default mode regions. Notably, HC subjects show higher time decay than patients with schizophrenia, and the apparent group differences present almost all over the brain. That also makes the data more useful for the classification. Intuitively, we feed subject-wise time decay data to multiple classifiers and get good accuracy in every case. The statelets are extracted from the whole dataset without using group (SZ/HC) information (unsupervised).

**Fig. 12:**
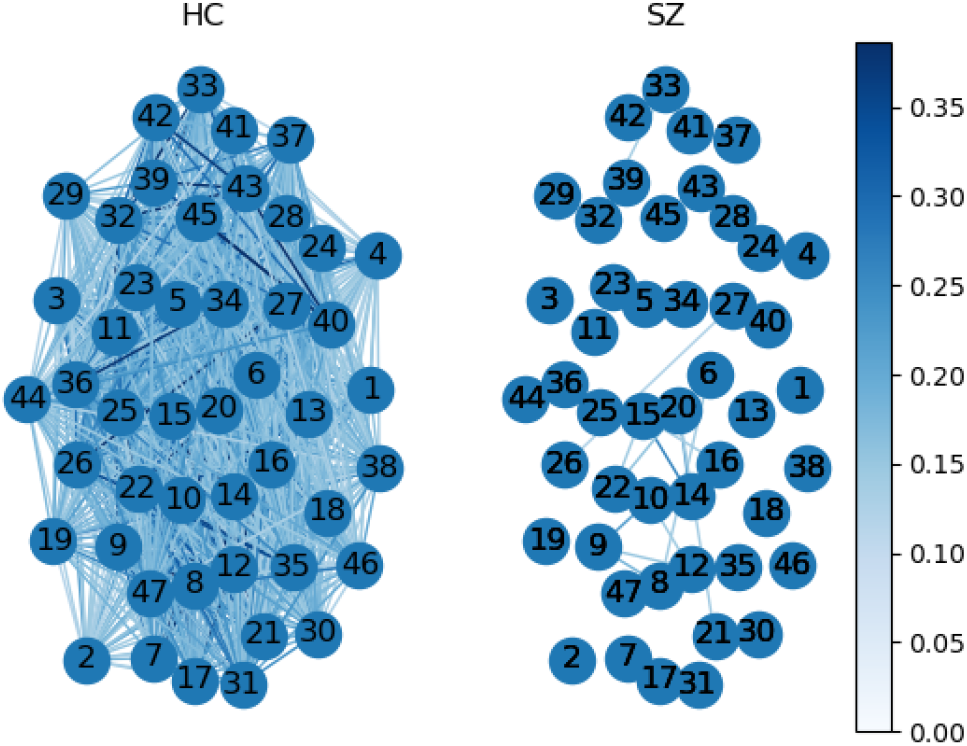
Time decay graphs from both groups. The nodes are functional networks, edges correspond to the connection between them (maximum 1081 possible), and the weight represents the mean time decay (TD) of a connection within a group dynamic. We compute color scaling from the 95 percentile of the total values. After thresholding at average group mean (SZ group mean + HC group mean)/2, 1061 edges survive in the HC group, and only 16 edges in the SZ groupFig. 13: Time decay graphs from both groups. The nodes are functional networks, edges correspond to the connection between them (maximum 1081 possible), and the weight represents the mean time decay (TD) of a connection within a group dynamic. We compute color scaling from the 95 percentile of the total values. After thresholding at average group mean (SZ group mean + HC group mean)/2, 1061 edges survive in the HC group and only 16 edges in the SZ group.

**Fig.13.**
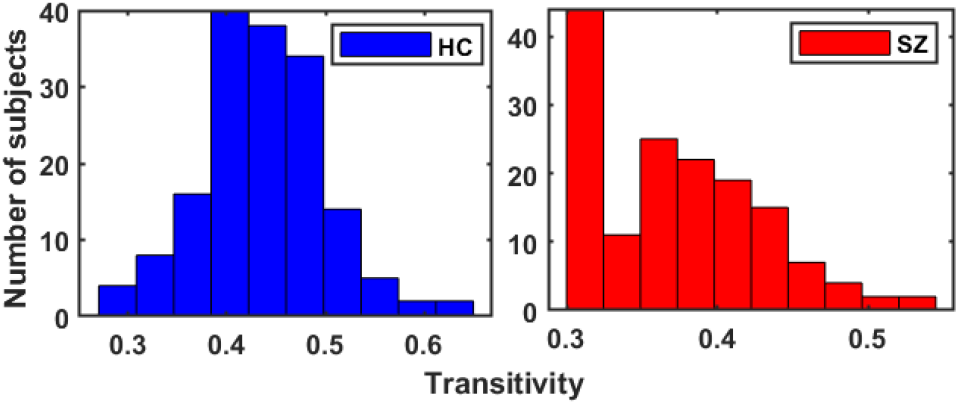
Histogram of transitivity from subject-level time decay graphs. It refers to the extent to which the relation between two nodes in a network connected by an edge is transitive. A significant portion of SZ subjects shows 0 transitivity, which means the connections are less consistent across different subjects. We show the differences are statistically significant using a two-sample t-test on both distributions. Transitivity is also related to the clustering coefficient.

**Fig. 14:**
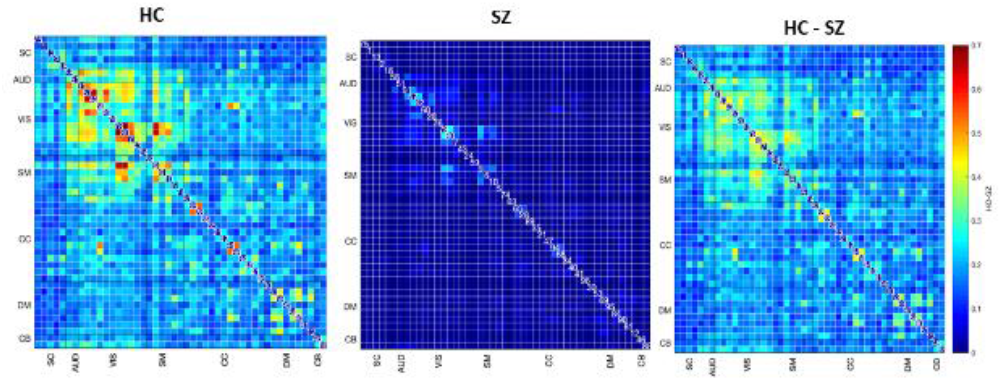
Pairwise time decays in patient and control groups and the HC - SZ group differences.

Thus, the classification accuracies are reasonably comparable. For clarity, we can consider it a dimensionality reduction method like principal component analysis (PCA) (Wold, Esbensen et al. 1987) for better features selection. We compare the performance between running the models directly on the dFNC matrix and the subject-wise time decay. For more robustness, we use multiple classifiers to test time decay and observe significant accuracy improvement. Figure 15 shows how the models perform in both cases. We can see a simple SVM or logistic regression model using time decay outperforms all other methods, including LSTM with attention.

**Fig. 15:**
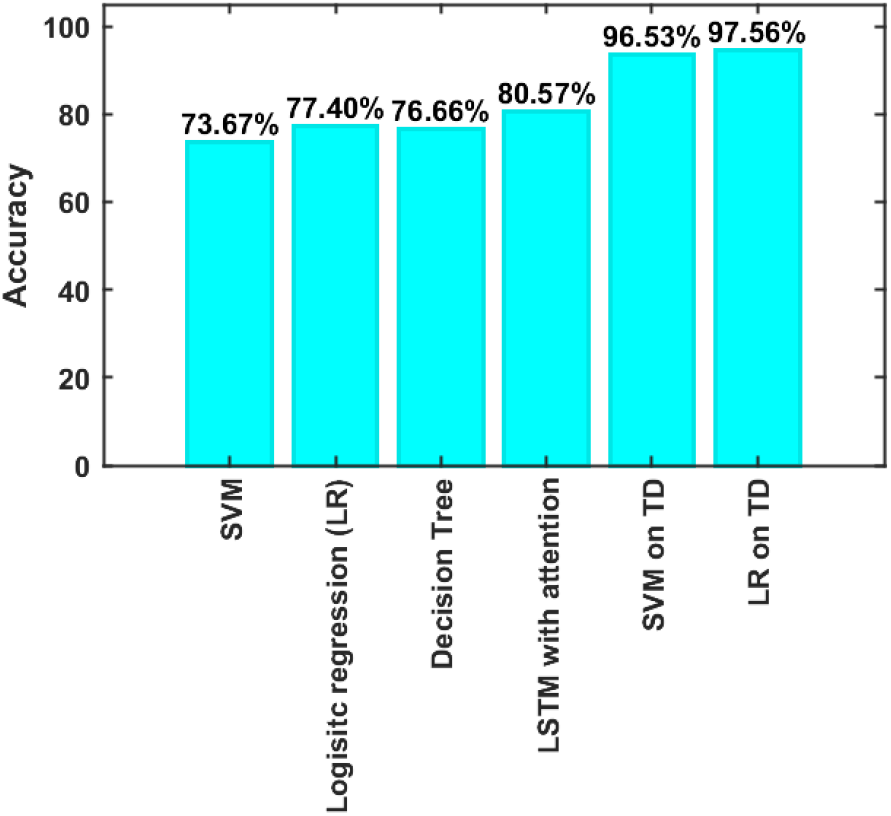
Classification accuracy for different methods. First, three methods were applied to the dFNC matrix and LSTM with the attention model applied to the dFNC time course. The last two methods use time decay (TD) for classification. We run the models on time decay information of all the subjects for 100 repeated iterations, and the accuracies are mean across the iterations. We train the model on 200 random samples in each iteration and cross-validate them using the remaining 114 subjects.

##### dFNC states from statelets

In another analysis, we focus on obtaining statelets across the dataset without separating SZ and HC subjects into two different bins. The idea is to extract the most frequent connectivity patterns from all the subjects, which is potentially extending the probability density of the statelets beyond the group dynamic. Here, we extract motifs from all the subjects and summarize them throughout the dataset instead of summarizing group-wise. The aim is to investigate how the motifs represent the subject’s dynamic irrespective of their group association and approximate statelets for all the data subgroups. Here, we consider all the subjects (SZ and HC) together in one dynamic to evaluate the statelets. We estimate the probability density profile for the motifs across all the subjects for a given connection. So, the density is no more group-specific rather throughout the dataset. We then run a tSNE on the collection of motifs extracted from the whole dataset; the global tSNE generates nine different statelets. Based on those statelets, we generate nine connectivity states and use the EMD of extracted motifs to determine their association towards a specific subgroup/state. That also determines the number of subjects within a state. Considering statelets as the centroid of the cluster, we assign all motifs to the nearest (lowest EMD) state. We evaluate the HC-SZ group differences in connectivity strength across the subjects within the state for each state. For measuring the statistical significance of these differences, we use a two-sample t-test. The t-values provide significant group differences in connectivity strength throughout the distinct regions of the brain (figure 16). HC subjects in state 1 show stronger connections than SZ, and the differences are statistically significant in sensorimotor (SM), cognitive control (CC), and default mode (DM) regions. In state 2 and 3, the differences are mostly HC < SZ; in state 3, group differences are statistically significant across the brain but similar strength in the connections between subcortical (SB) and other domains. State 4 shows mostly weaker differences except for a few components from auditory (AUD) and subcortical demonstrate SZ subjects are more strongly connected than HC in those regions. Two connections between AUD and DM show a statistically significant difference in state 5, and state 6 exhibits HC > SZ in the CC region. Above all, states 7, 8, and 9 show significant connectivity differences in CC and DM regions. Specifically, state 7 depicts HC > SZ and 8 & 9 shows HC < SZ group distinction. In an earlier study (Damaraju, Allen et al. 2014), strong group differences are mostly in VIS and AUD, but we got differences in SM, CC, and CB. We got both HC<SZ and SZ<HC in SM, CC, and DM. The most significant pattern of group differences we observe here is in CC, DM, and CB regions. This triangle shows the variation in connectivity between patients and controls across multiple states with an alternating directionality, which indicates a potential biomarker for specific subtypes of schizophrenia.

**Fig. 16.**
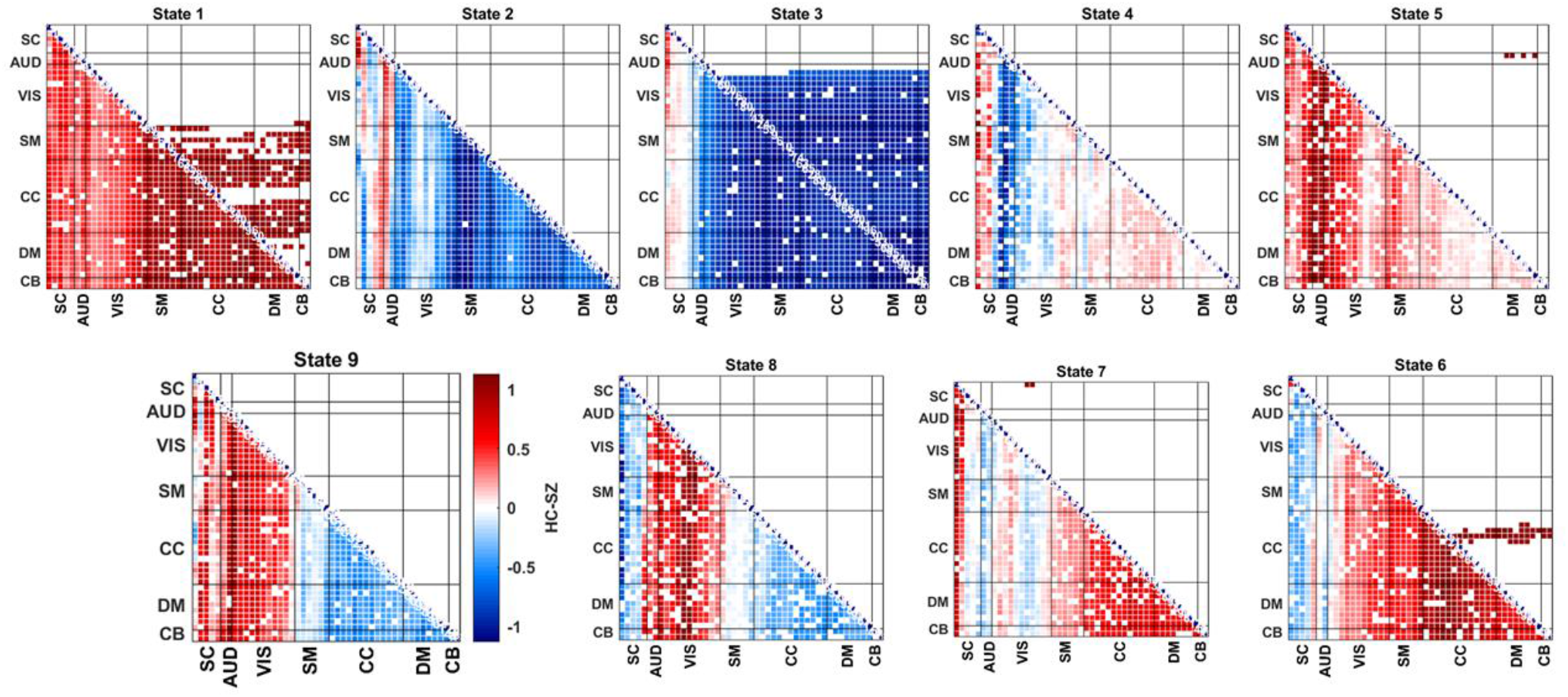
HC - SZ group differences in terms of max connectivity strength. State-wise group differences in functional connectivity (FC). We have both SZ and HC subjects’ groups at each state. For each pair of components, we have a subset of statelets from HC subjects and a subset from SZ subjects. Then, we compute the maximum connectivity strength of those statelets from both subgroups. A two-sample t-test using a null hypothesis of “No group difference” compares patients’ max connectivity vs. controls. A higher t-value indicates the rejection of the null hypothesis irrespective of their sign. However, the sign of t-values represents the directionality of the group difference. The pair matrix (47 × 47) is labeled into seven different domains subcortical (SB), auditory (AUD), visual (VIS), sensorimotor (SM), cognitive control (CC), default-mode (DM), cerebellar (CB), respectively. White cells in the matrix indicate either the absence of that pair or non-significant group differences for this pair within a state. The upper triangle represents the FDR corrected differences.

## 7. Conclusion

We proposed a novel method for analyzing dynamic functional connectivity via extracting high-frequency texture from the connectivity space. To our best knowledge, it is the first pattern mining application on dFNC data. The proposed framework addresses issues in current dFNC analysis methods by modeling the dynamics through brief shapes of the connectivity. The analysis of those motifs enables measuring the characteristics of brain circuitry and network organization. The major contributions are two-fold, an unsupervised method for motifs discovery using EMD as a distance metric and a probabilistic summarization of these patterns. Statelets exhibit handy outlines of the data that include abstraction, analytics, and current trends to the field-experts at a glance. Through statelets, we seek an improved understanding of connectivity states and the mechanism through which their dynamics vary across individuals. Results demonstrate how these state movies help to investigate the dynamic properties of an inherently dynamic system (i.e., brain). Our approach is more robust to noise while observing short-length dynamic features and ensures better interpretability in the results since the motifs itself describes the properties of functional association between a pair of networks. These connectivity shapes from SZ and HC dynamics help create a global contrast between healthy and diseased brains and illustrate crucial group differences in several dynamic properties like recurrence, modularity, synchronizability, etc. In a secondary analysis of dFNC states based on statelets, we observe distinctive group patterns in connectivity strengths. The difference in SM, CC, DM, and CB helps to highlight the dynamic disruptions in schizophrenia in those regions. Statelets enrich our understanding of how networks communicate in spontaneous brain activity and the pattern of binding/impairment between them to maximize information transfer efficiency and minimize connection costs. We believe this would fill in a gap in the field that will help us understand the dynamics better, possibly providing an improved way to study neuropsychiatric disorders more effectively.

## Author Contributions

Vince D. Calhoun (VDC) and Md Abdur Rahaman (MAR) proposed the idea of Statelets. MAR and Sergey M. Plis (SG) developed the methodology. MAR ran the experiments and drafted the manuscript. SG helped to design a few experiments and edited the manuscript. VDC supervised the whole project, thoroughly edited the paper, and gave valuable feedback on the results. Eswar Damaraju (ED) provided the data. ED and Debbrata Kumar Saha helped running the experiments and significantly contributed to the manuscript. All authors have approved the final version of the submission.

## Code and Data availability statement

The fBIRN dataset was collected ten years ago, and there is no data-sharing agreement created. Therefore, according to the IRB data privacy agreement, we are not allowed to share any subject-specific data. However, the resting-state fMRI data is available on request from the fBIRN project site; for the relevant paper, follow this link address https://www.ncbi.nlm.nih.gov/pmc/articles/PMC4651841/. The pre-processing pipelines are public, too, as referred to in the manuscript. All the pre-processing tools are available online at http://trendscenter.org/software.

